# Computer vision for pattern detection in chromosome contact maps

**DOI:** 10.1101/2020.03.08.981910

**Authors:** Cyril Matthey-Doret, Lyam Baudry, Axel Breuer, Rémi Montagne, Nadège Guiglielmoni, Vittore Scolari, Etienne Jean, Arnaud Campeas, Philippe Henri Chanut, Edgar Oriol, Adrien Meot, Laurent Politis, Antoine Vigouroux, Pierrick Moreau, Romain Koszul, Axel Cournac

**Affiliations:** Institut Pasteur, Unité Régulation Spatiale des Génomes, UMR 3525, CNRS, Paris, France; Sorbonne Université, Collège Doctoral, F-75005, Paris, France; Institut Pasteur, Computational Biology Department (CBD), Paris, France; ENGIE, Global Energy Management, Paris, France; Institut Pasteur, Synthetic Biology Laboratory, Paris, France

**Keywords:** Hi-C, loop calling, domain border, detection, quantification, Chromosight

## Abstract

Chromosomes of all species studied so far display a variety of higher order organizational features such as domains or loops often associated to biological functions and visible on Hi-C contact maps. We developed *Chromosight*, an algorithm inspired from computer vision that can detect patterns in Hi-C maps. *Chromosight* has greater sensitivity than existing methods, while being faster and applicable to any type of genomes, including bacteria, viruses, yeasts and mammals. Code and documentation: https://github.com/koszullab/chromosight

## Introduction

Proximity ligation derivatives of the chromosome conformation capture (3C) technique [1] such as Hi-C [2] or ChiA-PET [3] have unveiled a wide variety of chromatin 3D structures. These approaches determine the average contact frequencies between DNA segments within a genome, computed over hundreds of thousands of cells. In all species studied so far, sub-division of chromosomes into self-interacting domains associated with various functions have been observed. In addition, loops bridging distant loci within a chromosome (from a few kb to several Mb) are also commonly detected by Hi-C, such as during mammalian interphase [4] or yeast mitotic metaphase [5, 6].

Although most structural features can be identified by eye on the contact maps, automated detection is essential to quantify and facilitate the biological and physical interpretation of the data. While border detection can be achieved quite efficiently using different methods (segmentation, break-point detection, etc; [7]), loop calling remains challenging. Most tools aiming at detecting DNA loops in contact maps rely on statistical approaches and search for pixel regions enriched in contact counts, such as Cloops [8], HiCCUPS [9], HiCExplorer [10], diffHic [11], FitHiC2 [12], HOMER [13]. These programs can be computationally intensive and take several hours of computation for standard human Hi-C datasets (reviewed in [8]), or require specialized hardware such as GPU (HiCCUPS). In addition, most if not all of them were developed from and for human data. As a consequence, they suffer from a lack of sensitivity and fail to detect biologically relevant structures not only in non-model organisms (**Supplementary Fig. 3a**) but also in popular species such as yeast or bacteria. Besides this limitation, most programs are limited to domain or loop calling and remain unable to call *de novo* different contact patterns such as DNA hairpins or the asymmetric patterns seen in species such as Bacillus subtilis [14] (**Supplementary Fig. 1**).

Here we present *Chromosight*, an algorithm validated on mammalian, bacterial, viral and yeast genomes that detects and quantifies any type of pattern in chromosome contact maps, with a specific focus on chromosomal loops.

## Results

Chromosight takes a single, whole-genome contact map in sparse and compressed format as an input and processes each chromosome independently (balancing, detrending). A template representing a 3D structure (e.g. loop or border kernel) is fed to the program and sought for in the image of the contact map through two steps (**Fig. 1a**). First, the map is subdivided into sub-images correlated to the template; Then, sub-images with highest correlation values are labelled as template representations (methods). Correlation coefficients are computed by convolving the template over the contact map. To reduce computation time the template can be approximated using truncated singular value decomposition (tSVD) (Supplementary information). For selecting contiguous regions of high correlation values (foci), Chromosight uses Connected Component Labelling (CCL). Finally, the maximum is extracted within each focus and maps to the position of the detected pattern.

**Fig 1:**
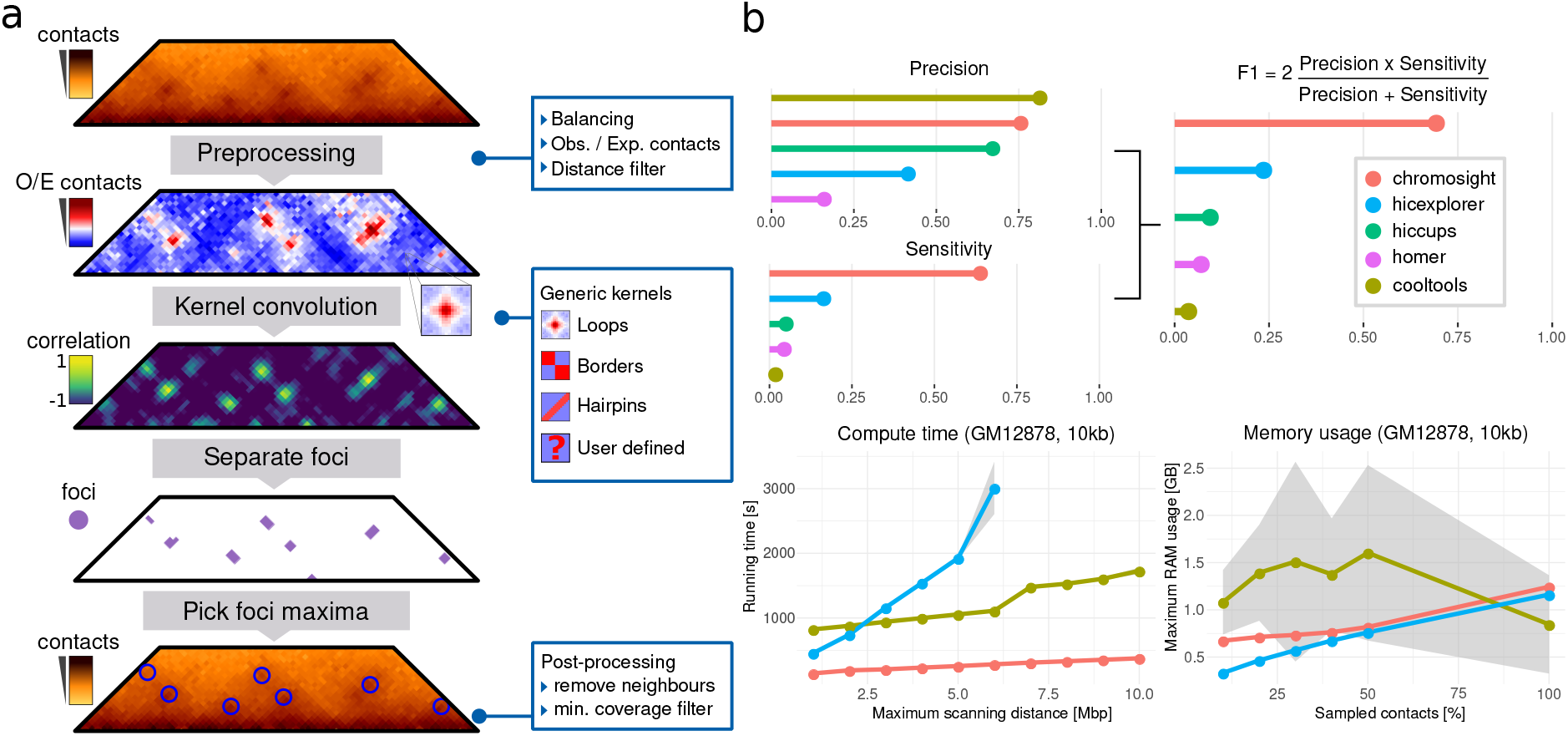
Chromosight workflow and benchmark. Matrix preprocessing involves normalization balancing followed by the computation of observed / expected contacts. Only contacts between bins separated by a user-defined maximum distance are considered. The preprocessed matrix is then convolved with a kernel representing the pattern of interest. For each pixel of the matrix, a Pearson correlation coefficient is computed between the kernel and the surrounding window. A threshold is applied on the coefficients and a connected component labelling algorithm is used to separate groups of pixels (i.e. foci) with high correlation values. For each focus, the coordinates with the highest correlation value is used as the pattern coordinates. Coordinates located in poorly covered regions are discarded. **b,** Comparison of Chromosight with different loop callers. Top: F1 score, Precision and Sensitivity scores assessed on labelled synthetic Hi-C data. Higher is better. **c,** Run-time **d,** Memory usage according to maximum scanning distance and the amount of downsampled contact events, respectively. Means and standard deviations (grey areas) are plotted.

We benchmarked Chromosight against 4 existing programs to judge the quality of loop calling performance on synthetic Hi-C data mimicking S. cerevisiae compact genome (methods; **Supplementary Fig. 1**). Whereas Chromosight displays a *precision* (proportion of true positives among detected patterns) comparable to the best programs, its *sensitivity* (proportion of relevant patterns detected) is nearly threefold higher ( 70% vs. 20% for the second best, Hicexplorer) (**Fig. 1b**). As a result, Chromosight F1 score, a metric that considers both precision and sensitivity, is also threefold higher, reflecting the effectiveness of the program at detecting more significant loops in compact genomes (**Supplementary Fig. 3a**). In addition Chromosight outperforms existing methods regarding computing time, running in a few minutes on high resolution human contact maps (**Fig. 1c**), without straining RAM (**Fig.1d**).

### Validation on experimental yeast data

Budding and fission yeast organize into chromatin loops during meiosis [15] and mitosis [5, 6, 16]. Recent work showed that S. cerevisiae mitotic loops are mediated by cohesins [5, 6]. Chromosight identified 974 loops along *S. cerevisiae* mitotic chromosomes [6] (**Fig. 2a**). An enrichment analysis shows that half (50%) of the anchors of mitotic loops are loci enriched in cohesin complex subunit Scc1 (**Fig. 2b**), (P < 10^−16^). The loop signal spectrum in mitosis shows the most stable loops are approximately 20 kb long (**Fig. 2c**). Interestingly, this size is also found in the *S.Pombe* yeast, which has longer chromosomes.

**Fig 2:**
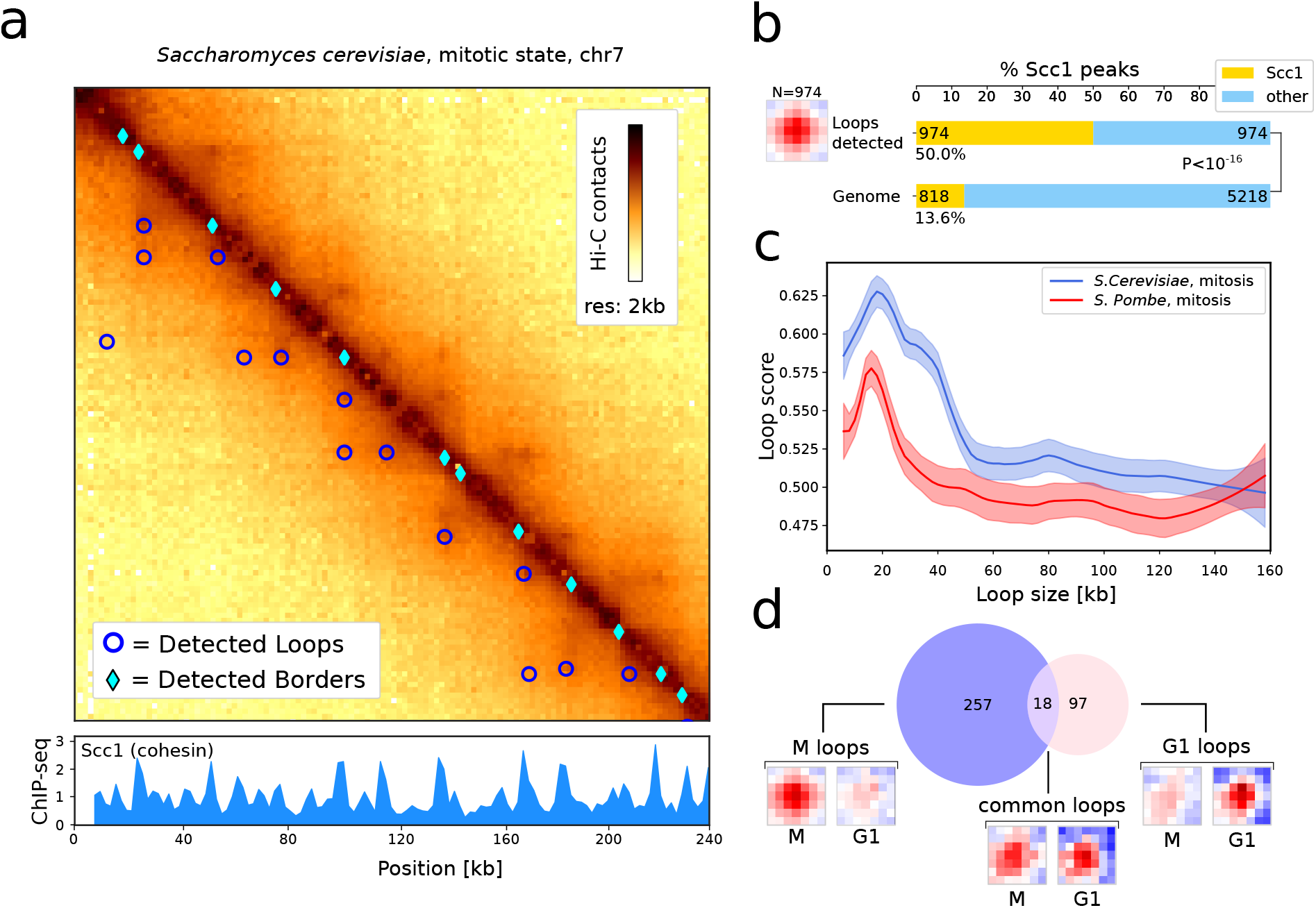
Applications on yeast genomes. **a,** Zoom-in of the contact map of chromosome 5 of *Saccharomyces cerevisiae* with synchronised ChIP-Seq signal of Scc1 protein (cohesin) at 2 kb resolution with detected loops and border patterns, [6]. **b,** Pileup plots of windows centered on detected loops with the number of detections. Barplots of the proportion of Scc1 peaks for detected loops and associated *p*-value (Fisher test). **c,** Loop spectrum showing scores in function of the loop size in *S. cerevisiae* and *S. Pombe*. Curves represent lowess-smoothed data for easier interpretation with confidence intervals. **d,** Number of loops detected only in G1 phase, M phase, or in both. For each category, the pileup of each set of coordinates is shown for both G1 and M conditions (sub-sampled with same number of contacts of G1).

On the other hand, loop calling on contact maps generated from cells lacking cohesin i.e in G1 still resulted in 115 loops, (**Fig. 2d**). Interestingly, this group of loops is different from the group of loops detected in the mitotic phase suggesting that cohesin independent processes act on chromosomal loop formation in yeast. Borders detection was also performed and shows that this pattern is associated with highly expressed genes (HEG) (**Supplementary Fig. 4**).

To validate the biological relevancy of the loops detected by the program during mitosis, we analyzed their dependency and association to cohesin using the quantification mode implemented in Chromosight (Methods, **Supplementary Fig. 5**).

### Exploration of various genomes

To further test the polyvalence of Chromosight, we called loops, borders and hairpins in Hi-C contact maps of human lymphoblastoids (GM12878) [17] (**Fig. 3a**). With default parameters, Chromosight identified 18,839 loops (compared to 10,000 elements previously detected [4]) whose basis fall mostly (≃58%, P < 10^−16^) into loci enriched in cohesin subunit Rad21, (**Fig. 3b**). Decreasing the detection threshold (Pearson coefficient parameter) allows to detect lower intensity but relevant patterns (**Supplementary Fig. 6**). The program also identified 9,638 borders, ≃75% of which coincide with CTCF binding sites, compared to ≃14% expected (P < 10^−16^). In human, self-interacting topologically associating domains (TADs) are delimited by CTCF-enriched sites, supporting that Chromosight correctly identifies domain boundaries. Finally, Chromosight detected 3,700 hairpin-like structures (**Fig. 3b**), a pattern not systematically sought for in Hi-C maps. Cohesin injection could account for the formation of such structures (**Supplementary Fig. 1b**). Such a mechanism is supported by the fact that chromosome coordinates for this detected group are enriched in cohesin loading factor NIPBL (2 fold effect, P < 10^−16^). The hairpin-like structures detected by Chromosight could therefore be interpreted as injection points for cohesin. To validate the results, we quantified loops and hairpins on contact maps generated from cells depleted either in cohesin or NIPBL, respectively which show a disappearance of the detected patterns (**Supplementary Fig. 7**). Finally, we detected the three patterns on various animals from the DNA Zoo project [18], showing that stable loops of ≃100 kb is a conserved feature of animal genomes (**Supplementary Fig. 8**).

**Fig 3:**
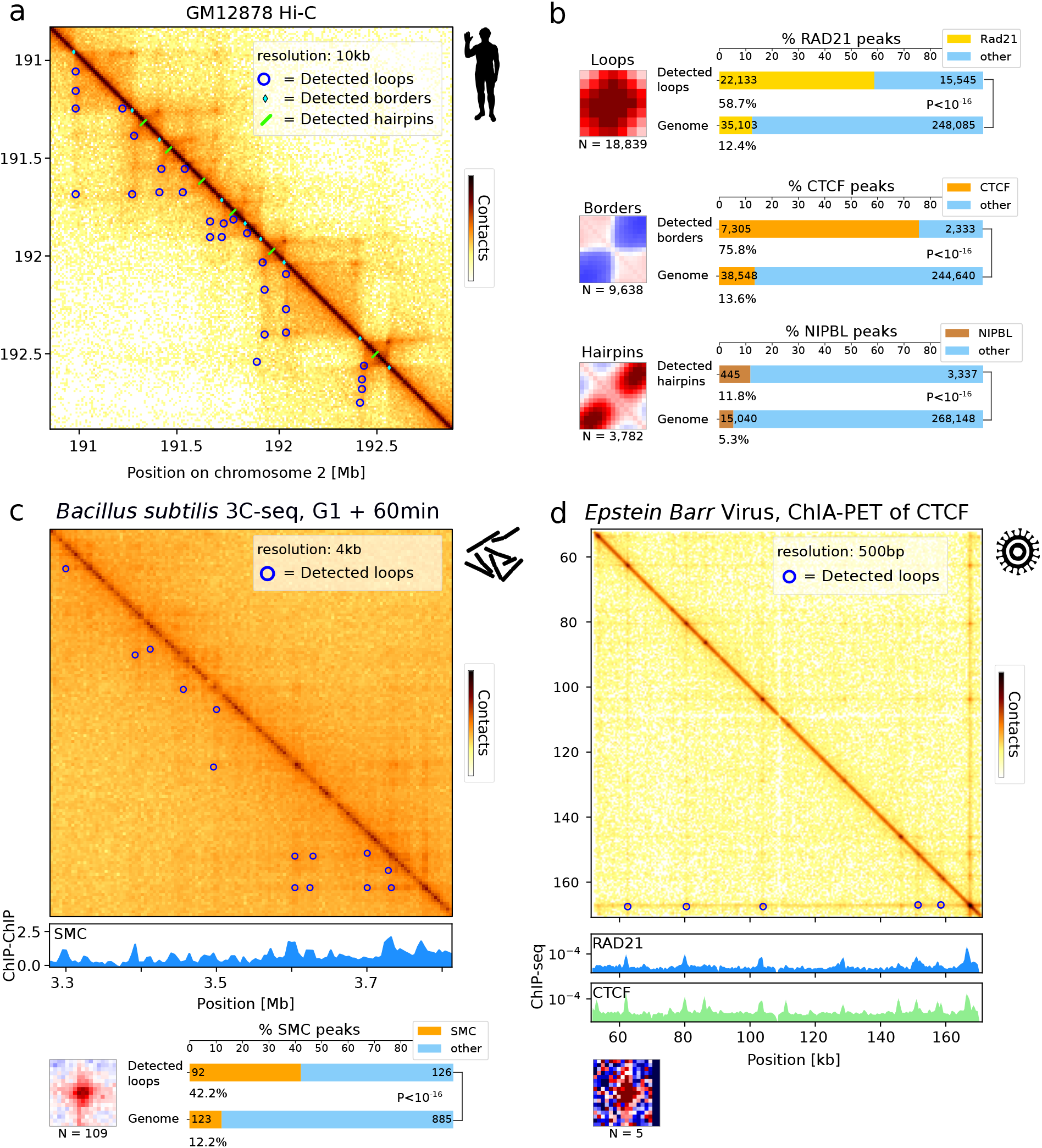
Applications to various genomes. **a,** Zoom of contact map for chromosome 2 of *Homo Sapiens* at 10 kb resolution [17] with Chromosight detection of loop, border and hairpin patterns. **b,** pileup plots of windows centered on detected loops, borders and hairpins with the number of detections. Bar plots showing proportion in Rad21 peaks for detected loops, proportion in CTCF peaks for detected borders and proportion of NIPBL peaks for detected hairpins and associated pvalue (Fisher test). **c,** Detection of loops in *Bacillus sutilis* genome. Zoom of contact map with annotated detected loops synchronised with CHIP-chip signal of SMC [14]. Associated pileup plot of the detections and bar plot showing enrichment of SMC in the anchors of the detected loops (Fisher test). **d,** Dectection of loops in Epstein Barr Virus genome, [3] synchronized with ChIP-seq signal of Rad21 and CTCF. Associated pileup plot of the detections.

The loop detection efficiency was also tested using genomic contact maps of the bacteria *Bacillus subtilis* [14]. Despite the relatively noisy 3Cseq data, Chromosight identified 109 loops distributed throughout the chromosome (**Fig. 3c**), including large ones bridging loci separated by more than 100kb (**Supplementary Fig. 9**). Annotation of loop anchor positions showed a strong enrichment with the SMC complex condensin (**Fig. 3c**). The detection of loops at sizes larger than 100 kb is surprising in comparison to the sizes detected in yeast (20 kb) and humans (130 kb).

Finally, we used Chromosight to detect loops on contact data generated using paired-end tag sequencing (ChIA-PET) [3], which captures contacts between DNA segments associated to a protein of interest. Surprisingly, Chromosight detected several loops (5) inside the genome of the Epstein Barr virus [3]. These loops, of a few dozen kb in size, coincide with the position of the cohesin (Rad21) and CTCF binding sites present along the virus genome (**Fig. 3d**). Such interactions have been suggested from 3C qPCR data [19]. Automatic detection now unambiguously supports a novel viral chromosome structure that could impact the transcriptional regulation and metabolism of the virus [19].

## Discussion

We demonstrated that Chromosight outperforms other loop detection methods, and that it can be used to extract other biologically relevant patterns from various contact technologies. We expect that additional pattern configurations will be added by the community, such as stripes, bow patterns, patterns associated to misassemblies or structural variations (e.g. inversion, translocations) or any pattern of interest that the user can select directly when viewing the data, since a ‘Click and find’ mode was included for that purpose in the program (**Supplementary Fig. 10**).

With improved sequencing costs, new designs of experimental protocols and methods for amplifying specific genomic regions, contact data should achieve higher resolutions and reveal new spatial structures. The algorithmic approach we presented here provides a computational and statistical framework for the discovery of new principles governing chromosome architecture.

## Supporting information

Supplementary informations

## Acknowledgements

This work was initiated during a Hackathon between Institut Pasteur scientists and ENGIE engineers. We would like to thank all the people that allow the organization of this event especially Anne-Gaelle Coutris, Romain Tchertchian and Olivier Gascuel. Frédéric Beckouët and all the members of Spatial Regulation of Genomes unit are thanked for stimulating discussions and feedbacks. This work used the computational and storage services (TARS cluster) provided by the IT department at Institut Pasteur, Paris. C.M-D was supported by the Pasteur - Paris University (PPU) International PhD Program. A.B. works within the framework of a “Mécénat Compétence” contract of the company ENGIE. This research was supported by funding to R.K. from the European Research Council under the Horizon 2020 Program (ERC grant agreement 260822) and by ANR JCJC 2019, “Apollo” allocated to A.C.

## Author contributions

All authors contribute to the design of the algorithm. CMD, AB, LB, AC implement it. CMD, RM, LB compare to other algorithms. LB and AC design strategy for simulations of data. CMD, PM, RK and AC analyzed biological data and interpret results. CMD, AB, LB, RK and AC prepared the manuscript. All authors read and approved the final manuscript.

## Data availability

Benchmark data available on zenodo (doi: 10.5281/zenodo.3742095). All other data associated with this study are available from the corresponding authors.

## Code availability

Software and documentation available at https://github.com/koszullab/chromosight.

## Methods

### Simulation of Hi-C matrices

Simulated matrices were generated using a bootstrap strategy based on Hi-C data from mitotic *S. cerevisiae* [20] at 2 kb resolution. Three main features were extracted from the yeast contact data (**Supplementary Fig. 2**): the probability of contact as a function of the genomic distance (*P* (*s*)), the positions of borders detected by HicSeg [21] and positions of loops detected manually on chromosome 5. Positions from loops and borders were then aggregated into pileups of 17×17 pixels. We generated 2000 simulated matrices of 289×289 pixels. A first probability map of the same dimension is generated by making a diagonal gradient from *P* (*s*) representing the polymer behaviour. For each of the 2000 generated matrices, two additional probability maps are generated. The first by placing several occurrences of the border pileup on the diagonal, where the distance between borders follows a normal distribution fitted on the experimental coordinates. The second probability map is generated by adding the loop kernel 2-100 pixels away from the diagonal with the constraint that it must be aligned vertically and horizontally with border coordinates. For each generated matrix, the product of the P(s), borders and loops probability maps is then computed and used as a probability law to sample contact positions while keeping the same number of reads as the experimental map.

### Benchmarking

To benchmark precision, sensitivity and F1 score, the simulated Hi-C data set with known loop coordinates were used. Each algorithm was run with a range of 60-180 parameter combinations (**Supplementary Fig. 3**) on 2000 simulated matrices and F1 score was calculated on the ensemble of results for each parameter combination separately (**Supplementary Table 1**). For each software, scores used in the final benchmark (**Fig. 1**) are those from the parameter combination that yielded the highest F1 score.

For the performance benchmark, HiCCUPS and HOMER were excluded. The former because it runs on GPU, and the latter because it uses genomic alignments as input and is much slower. The dataset used is a published high coverage Hi-C library [22] from human lymphoblastoid cell lines (GM12878). To compare RAM usage across programs, this dataset was downsampled at 10, 20, 30, 40 and 50% contacts and the maximum scanning distance was set to 2 Mbp. To compare CPU time, all programs were run on the full dataset, at different maximum scanning distances, with a minimum scanning distance of 0 and all other parameters left to default. All programs were run on a single thread, on a Intel(R) Core(TM) i7-8700K CPU at 3.70GHz with 32GB of available RAM.

Software versions used in the benchmark are Chromosight v0.9.0, hicexplorer v3.3.1, cooltools v0.2.0, homer 4.10 and hiccups 1.6.2. Input data, scripts and results of both benchmarks are available on zenodo (doi: 10.5281/zenodo.3742095)

### Preprocessing of Hi-C matrices

Chromosight accepts input Hi-C data in cool format [23]. Prior to detection, Chromosight balances the whole genome matrix using the ICE algorithm [24] to account for Hi-C associated biases. For each intrachromosomal matrix, the observed/expected contact ratios are then computed by dividing each pixel by the mean of its diagonal. This erases the diagonal gradient due to the power-law relationship between genomic distance and contact probability, thus emphasizing local variations in the signal (**Fig. 1a**). Intra-chromosomal contacts above a user-defined distance are discarded to constrain the analysis to relevant scales and improve performances.

### Calculation of Pearson coefficients

Correlation coefficients are computed by convolving the template over the contact map. Convolution algorithms are often used in computer vision where images are typically dense. Hi-C contact maps, on the other hand, can be very sparse. Chromosight’s convolution algorithm is therefore designed to be fast and memory efficient on sparse matrices. It can also exclude missing bins when computing correlation coefficients. Those bins appear as white lines on Hi-C matrices and can be caused by repeated sequences or low coverage regions.

The contact map can be considered an image IMG_CONT_ where the intensity of each pixel img_cont_[*i, j*] represents the contact probability between loci *i* and *j* of the chromosome. In that context, each pattern of interest can be considered a template image IMG_TMP_ with *M*_tmp_ rows and *N*_TMP_ columns.

The correlation operation consists in sliding the template (IMG_TMP_) over the image (IMG_CONT_) and measuring, for each template position, the similarity between the template and its overlap in the image. We used the Pearson correlation coefficient as a the measure of similarity between the two images. The output of this matching procedure is an image of correlation coefficients IMG_CORR_ such that

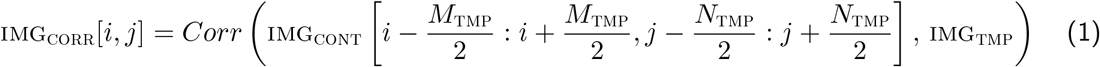

where the correlation operator *Corr*(*·, ·*) is defined as

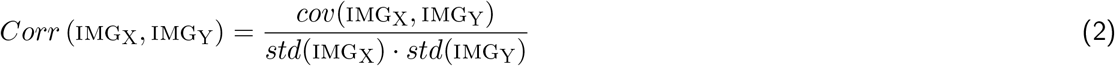

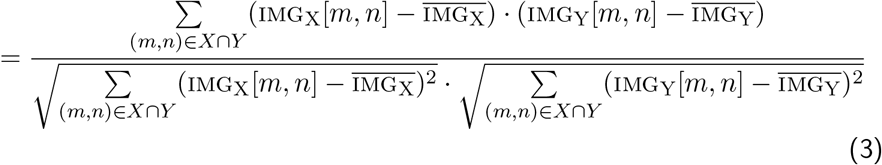

where 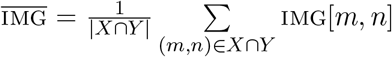 IMG[*m, n*], *X* ∩ *Y* is the set of pixel coordinates that are *valid* in image IMG_X_ and in image IMG_Y_, and |*X* ∩ *Y*| is the number of *valid* pixels in IMG_X_ and IMG_Y_. A pixel in IMG_CONT_ is defined as *valid* when it is outside a region with missing bins.

### Separation of high-correlation foci

Selection is done by localizing specific local maxima within IMG_CORR_. We proceeded as follows: first, we discard all points (*i, j*) where IMG_CORR_[*i, j*] *τ*_CORR_. An adjacency graph *A_dxd_* is then generated from the *d* remaining points. The value of *A*[*i, j*] is a boolean indicating the (4-way) adjacency status between the *i^th^* and *j^th^* nonzero pixels. The scipy implementation of the CCL algorithm for sparse graphs [25] is then used on *D* to label the different contiguous foci of nonzero pixels. Foci with less than two pixels are discarded. For each focus, the pixel with the highest coefficient is determined as the pattern coordinate.

Patterns are then filtered out if they overlap too many empty pixels or are too close from another detected pattern. The remaining candidates in IMG_CORR_ are scanned by decreasing order of magnitude: every time a candidate is appended to the list of selected local maxima, all its neighboring candidates are discarded. The proportion of empty pixels allowed and the minimum separation between two patterns are also user defined parameters.

### Biological analyses

Pairs of reads were aligned independently using Bowtie2 (v2.3.4.1) with *–very-sensitive-local* against the *S. cerevisiae* SC288 reference genome (GCF000146045.2). Uncuts, loops and religation events were filtered as described in [26]. Contact data were binned at 2 kb and normalised using the ICE balancing method [24]. The detection on yeast data was performed with default parameters using a 7×7 loop kernel available in Chromosight using *–pattern loops small* unless mentioned otherwise. For enrichment analysis, cohesin peaks were defined using ChIP-seq data from [27]. Raw reads were aligned with bowtie2 and only mapped positions with Mapping Quality superior to 30 were kept and signals were also binned at 2 kb to synchronise with Hi-C data. Peaks of cohesins were considered with ChIP / input > 1.5 and peaks closer than 10kb to centromeres or rDNA were removed.

Annotation of highly expressed genes was done using RNA-seq data from [28]. Alignment was done as above. The distribution of the number of reads for each 2kb bin was computed and the top 20% of the distribution were considered bins with high transcription. For border annotation, a set of plus or minus 1 bin on the detected positions is used. For human data, hg19 genome assembly was used with same strategy for alignment, construction and normalisation of contact data. ChIPseq peaks were retrieved from UCSC database (**Supplementary Table 2**). *Bacillus subtilis* data were aligned with the PY79 genome version and the SMC signal was extracted using ChIP-chip data from [29] and processed as described previously [30]. Peaks were annotated with the *find peaks* function from scipy (v1.4.1), with parameters *threshold = 0.1, width = 50*. CHiA-PET data were processed as Hi-C data except that the contact maps were binned at a 500bp resolution. Epstein-Barr virus (EBV) genome, strain B95-8 (V01555.2) sequence was used to align the reads from EBV.

