## Supplementary informations for "Computer vision for pattern detection in chromosome contact maps"

### Fast 2D-convolution via SVD

Optionally, Chromosight's convolution algorithm can be accelerated further by approximating the template. This is done using truncated singular value decomposition (tSVD) to decompose the template into two sets of vectors whose product contain most of the information in the template, while reducing the number of operations needed in the convolution. This note explains the acceleration of 2D-convolution by using the SVD decomposition of the kernel. It was inspired from section 6.4.2 in *Computer and Robot Vision Vol. I* by Haralick and Shapiro (1992) [1].

#### The general case

Suppose that the contact map $\mathrm{IMG}_{\mathrm{CONT}}$ and the template $\mathrm{IMG}_{\mathrm{TMP}}$ have respectively size $(M_{\mathrm{CONT}}, N_{\mathrm{CONT}})$ and $(M_{\mathrm{TMP}}, N_{\mathrm{TMP}})$.

The convolution of $\mathrm{IMG}_{\mathrm{CONT}}$ by $\mathrm{IMG}_{\mathrm{TMP}}$, noted $\mathrm{IMG}_{\mathrm{CONT}} * \mathrm{IMG}_{\mathrm{TMP}}$, is an array such that

$$\left(\mathrm{IMG}_{\mathrm{CONT}} * \mathrm{IMG}_{\mathrm{TMP}}\right)[i,j] := \sum_{m=0}^{M_{\mathrm{TMP}}-1} \sum_{n=0}^{N_{\mathrm{TMP}}-1} \mathrm{IMG}_{\mathrm{CONT}}\left[i+m, j+n\right] \times \mathrm{IMG}_{\mathrm{TMP}}[m,n] \tag{1}$$

for $i = 1, \ldots, M_{\mathrm{CONT}} - M_{\mathrm{TMP}} + 1$ and $j = 1, \ldots, N_{\mathrm{CONT}} - N_{\mathrm{TMP}} + 1$. Otherwise stated $\mathrm{IMG}_{\mathrm{CONT}*\mathrm{TMP}} := \mathrm{IMG}_{\mathrm{CONT}} * \mathrm{IMG}_{\mathrm{TMP}}$ is an array of size $(M_{\mathrm{CONT}} - M_{\mathrm{TMP}} + 1, N_{\mathrm{CONT}} - N_{\mathrm{TMP}} + 1)$.

The computation of $\left(\mathrm{IMG}_{\mathrm{CONT}} * \mathrm{IMG}_{\mathrm{TMP}}\right)[i,j]$ requires $2\,M_{\mathrm{TMP}}\,N_{\mathrm{TMP}}$ operations, composed of $M_{\mathrm{TMP}}\,N_{\mathrm{TMP}}$ additions and $M_{\mathrm{TMP}}\,N_{\mathrm{TMP}}$ multiplications.

#### The separable case

Suppose that the template is *separable* i.e. there exists two vectors $\mathrm{U}_{\mathrm{TMP}}$ and $\mathrm{V}_{\mathrm{TMP}}$, with respective size $(M_{\mathrm{TMP}})$ and $(N_{\mathrm{TMP}})$, such that

$$\mathrm{IMG}_{\mathrm{TMP}}[m,n] = \mathrm{U}_{\mathrm{TMP}}[m]\,\mathrm{V}_{\mathrm{TMP}}[n]. \tag{2}$$

The operations in equation (1) can then be re-arranged more efficiently:

$$\begin{aligned}\left(\mathrm{IMG}_{\mathrm{CONT}} * \mathrm{IMG}_{\mathrm{TMP}}\right)[i,j] &= \sum_{m=0}^{M_{\mathrm{TMP}}-1} \left( \sum_{n=0}^{N_{\mathrm{TMP}}-1} \mathrm{IMG}_{\mathrm{CONT}}\left[i+m, j+n\right] \times \mathrm{V}_{\mathrm{TMP}}[n] \right) \times \mathrm{U}_{\mathrm{TMP}}[m] \qquad (3)\\ &= \sum_{m=0}^{M_{\mathrm{TMP}}-1} \mathrm{IMG}_{\mathrm{CONT}*\mathrm{V}_{\mathrm{TMP}}}[i+m, j] \times \mathrm{U}_{\mathrm{TMP}}[m] \qquad (4)\end{aligned}$$

where

$$\mathrm{IMG}_{\mathrm{CONT}*\mathrm{V}_{\mathrm{TMP}}}[i,j] := \sum_{n=0}^{N_{\mathrm{TMP}}-1} \mathrm{IMG}_{\mathrm{CONT}}\left[i+m, j+n\right] \times \mathrm{V}_{\mathrm{TMP}}[n] \tag{5}$$

The computation of an element $\mathrm{IMG}_{\mathrm{CONT}*\mathrm{TMP}}[i,j]$ costs $N_{\mathrm{TMP}}$ multiplications and $N_{\mathrm{TMP}}$ additions. According to equation (4), the computation of $\mathrm{IMG}_{\mathrm{CONT}*\mathrm{V}_{\mathrm{TMP}}}[i,j]$ requires $M_{\mathrm{TMP}} + N_{\mathrm{TMP}}$ multiplications and $M_{\mathrm{TMP}} + N_{\mathrm{TMP}}$ additions.

Consequently, the evaluation of $\text{IMG}_{\text{CONT}*\text{TMP}}[i,j]$ costs $2\,(M_{\text{TMP}} + N_{\text{TMP}})$ operations in the separable case, which compares favorably to the $2\,M_{\text{TMP}}\,N_{\text{TMP}}$ operations required in the general case.

**The SVD case**

Next, suppose that the template has a representation as the sum of $K$ separable kernels:

$$\text{IMG}_{\text{TMP}}[m,n] = \sum_{k=1}^{K} \text{U}_{\text{TMP}}[m,k]\,\text{V}_{\text{TMP}}[k,n]. \tag{6}$$

The number of operations involved in evaluating $\text{IMG}_{\text{CONT}*\text{TMP}}[i,j]$ is $2\,(M_{\text{TMP}} + N_{\text{TMP}})$ for each kernel plus $K-1$ additions necessary to sum up the contribution of each kernel. In total, there are hence $2\,K\,(M_{\text{TMP}} + N_{\text{TMP}}) + K - 1$ operations.

The template $\text{IMG}_{\text{TMP}}$ is not necessarily equal to the superposition of $K$ separable kernels, but it can always be approximated by such a superposition. The (truncated) SVD algorithm discussed below allows to construct such an approximation.

The Singular Value Decomposition (SVD) factorizes any rectangular matrix $A$ of size $(M,N)$ as

$$A = U\,D\,V \tag{7}$$

where $U$ is a $(M,M)$ orthogonal matrix, V is a $(N,N)$ orthogonal matrix and $D$ is a $(M,N)$ matrix all of whose nonzero entries are on the diagonal and are positive.

Given any template $\text{IMG}_{\text{TMP}}$, it is possible to approximate it by retaining only the $K$ largest singular values in the SVD of $A = \text{IMG}_{\text{TMP}}$, such that:

$$\text{U}_{\text{TMP}}[:,k] = \sqrt{D[k,k]}\,U[:,k] \tag{8}$$

$$\text{V}_{\text{TMP}}[k,:] = \sqrt{D[k,k]}\,V[k,:] \tag{9}$$

$$\tag{10}$$

Let's give a toy example of the operations spared by using a SVD approach. Suppose that $M_{\text{TMP}} = N_{\text{TMP}} = 17$, the brute force convolution would require $2 \times 17 \times 17 = 578$ operations per point. In contrast, if we use a SVD convolution with $K=1$, the number of operations reduces to $68$, which represents only $12\%$ of the brute force approach. Even with $K=8$, we are below $50\%$ of the brute force approach.

### Additional Figures

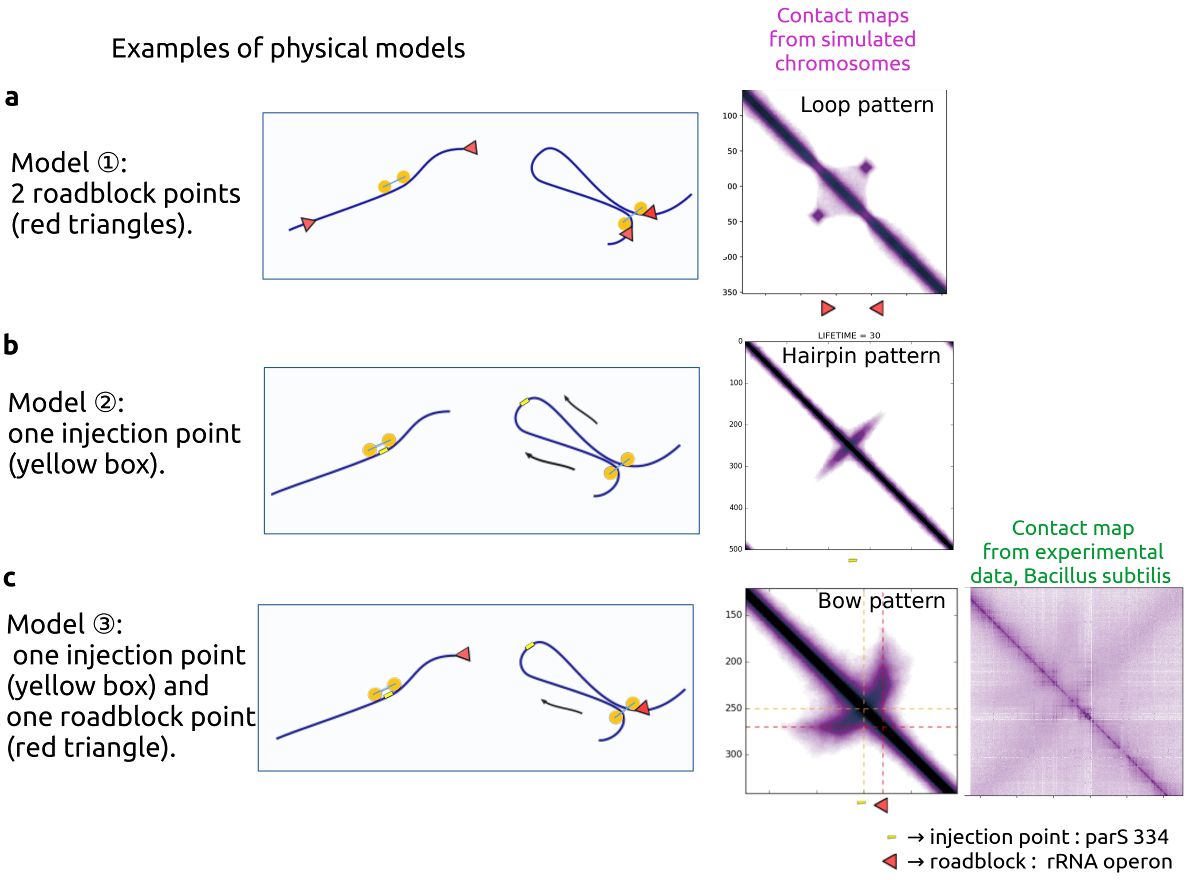


**Fig. 1: Toy models that can link visual patterns and physical models. a,** Model with loop extruding motors and two roadblock points leading to loop pattern. **b,** Model with loop extruding motors and one injection point leading to hairpin pattern. **c,** Model with loop extruding motors, one injection point and one roadblock leading to bow pattern. This pattern have been observed in contact data from Bacillus subtilis bacteria [2]. By connecting the simulation and experimental contact data, the identified roadblock is a highly transcribed gene, (rDNA operon) and the injection site corresponds to the ParS 334 site. Molecular dynamics simulations were performed using OpenMM [3] and libraries with default parameters of [4].

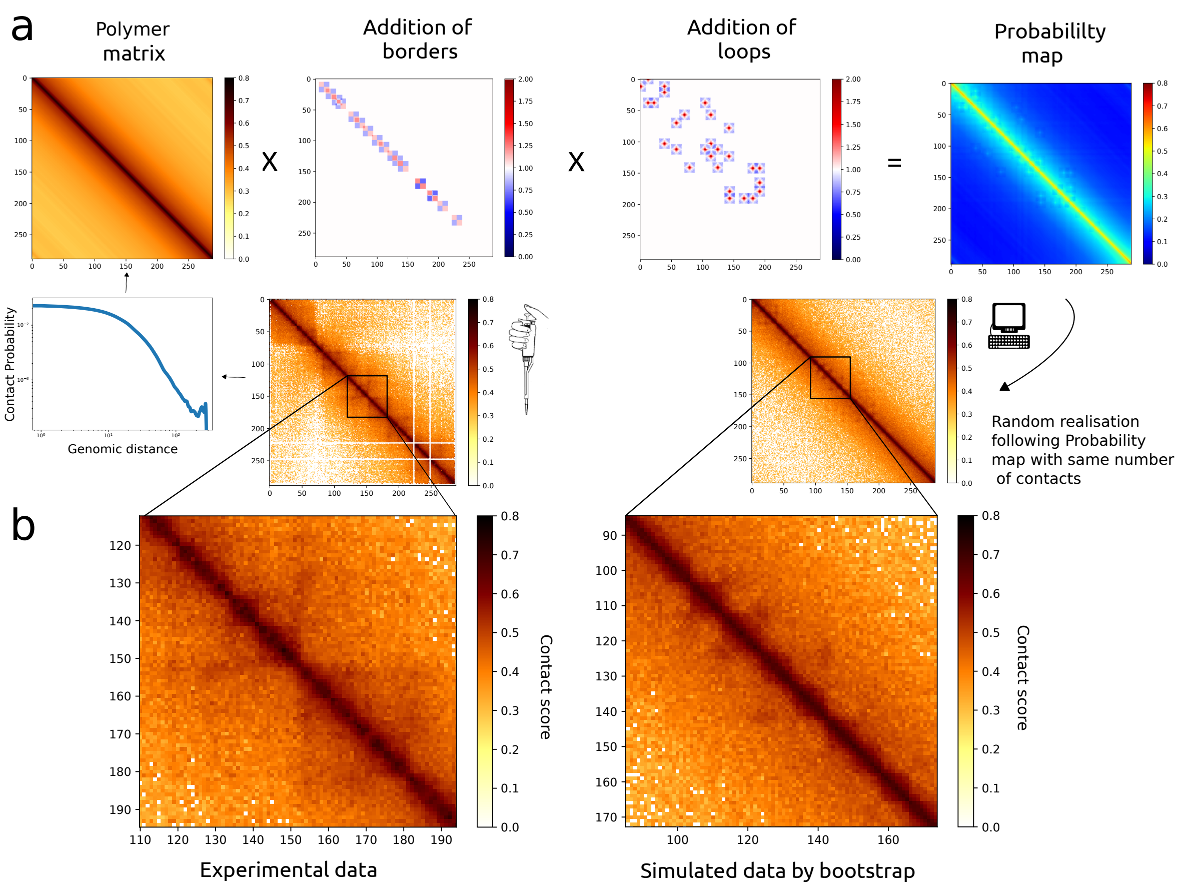


**Fig. 2: Strategy for the generation of simulated contact data for benchmark tests of different algorithms of loop calling a,** The simulated data were generated with a boostrap approach based on contact data generated for yeast *S.Cerevisiae* in mitotic phase [5]. Three main features of the contact data were extracted: the probability of contact as a function of the genomic distance (P(s), Polymer matrix), presence of borders, presence of loops. The positions and intensity of border and loop patterns were defined thanks to pile-up signals from patterns detected by eye on the contact maps. Their positions were chosen according to a law of probabilities based on experimental data (see Methods). The product of the 3 feature matrices results in a probability matrix (**a**, right). This matrix is used as a probability law to select contact positions while keeping the same number of reads as the experimental map. **b,** Zoom of contact maps for experimental and simulated data showing patterns of loops and borders.

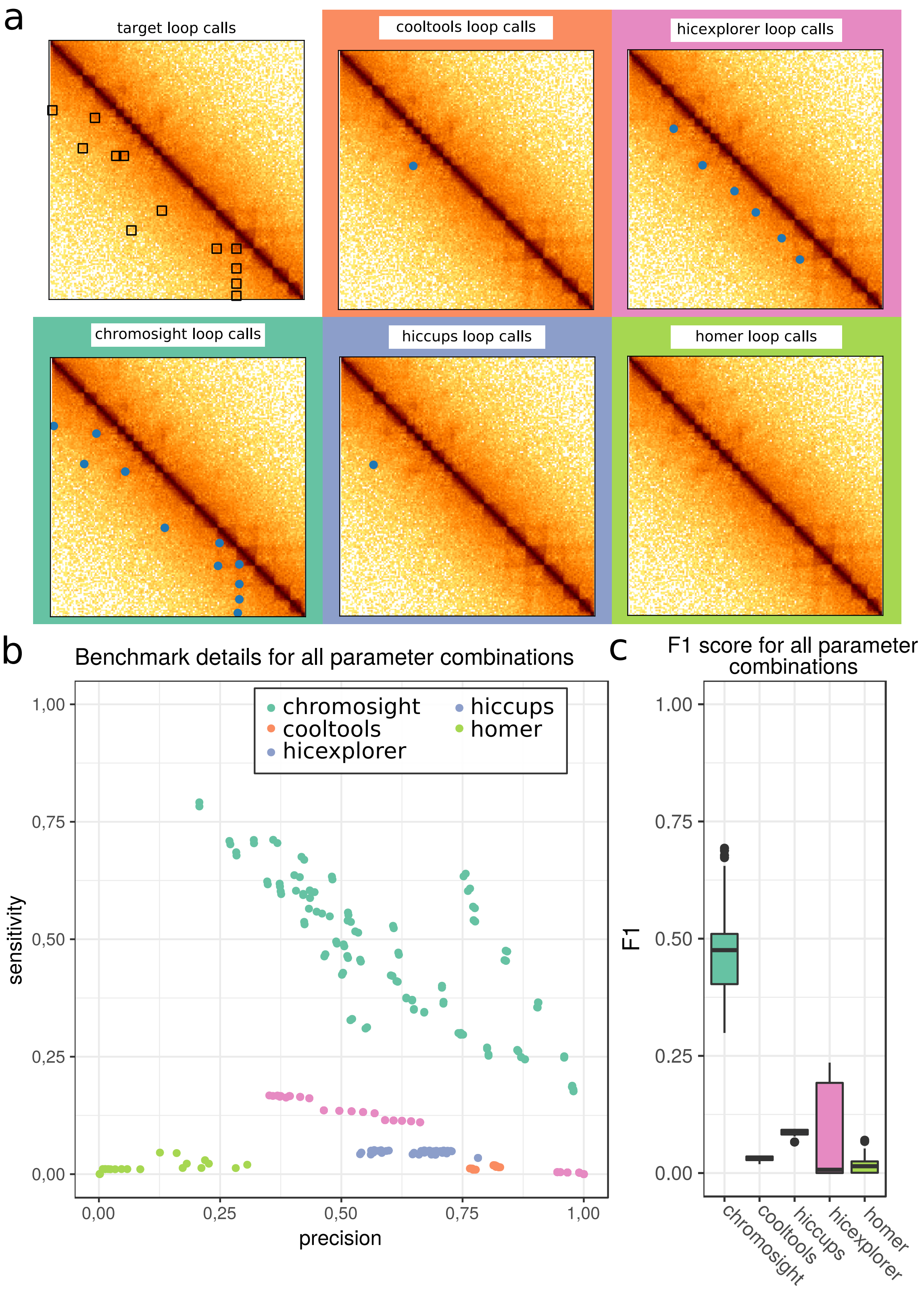


**Fig. 3: Comparison of different loop callers** Detailed benchmark results. a) Example region from a synthetic matrix with real loop calls (top left) and loops detected by all algorithms used in the benchmark using the combinations of parameters which yielded the highest F1 score. b) Precision and sensitivity from all algorithms on synthetic matrices, on the whole range of parameters tested. c) Distribution of F1 Scores for each algorithm for the range of parameters.

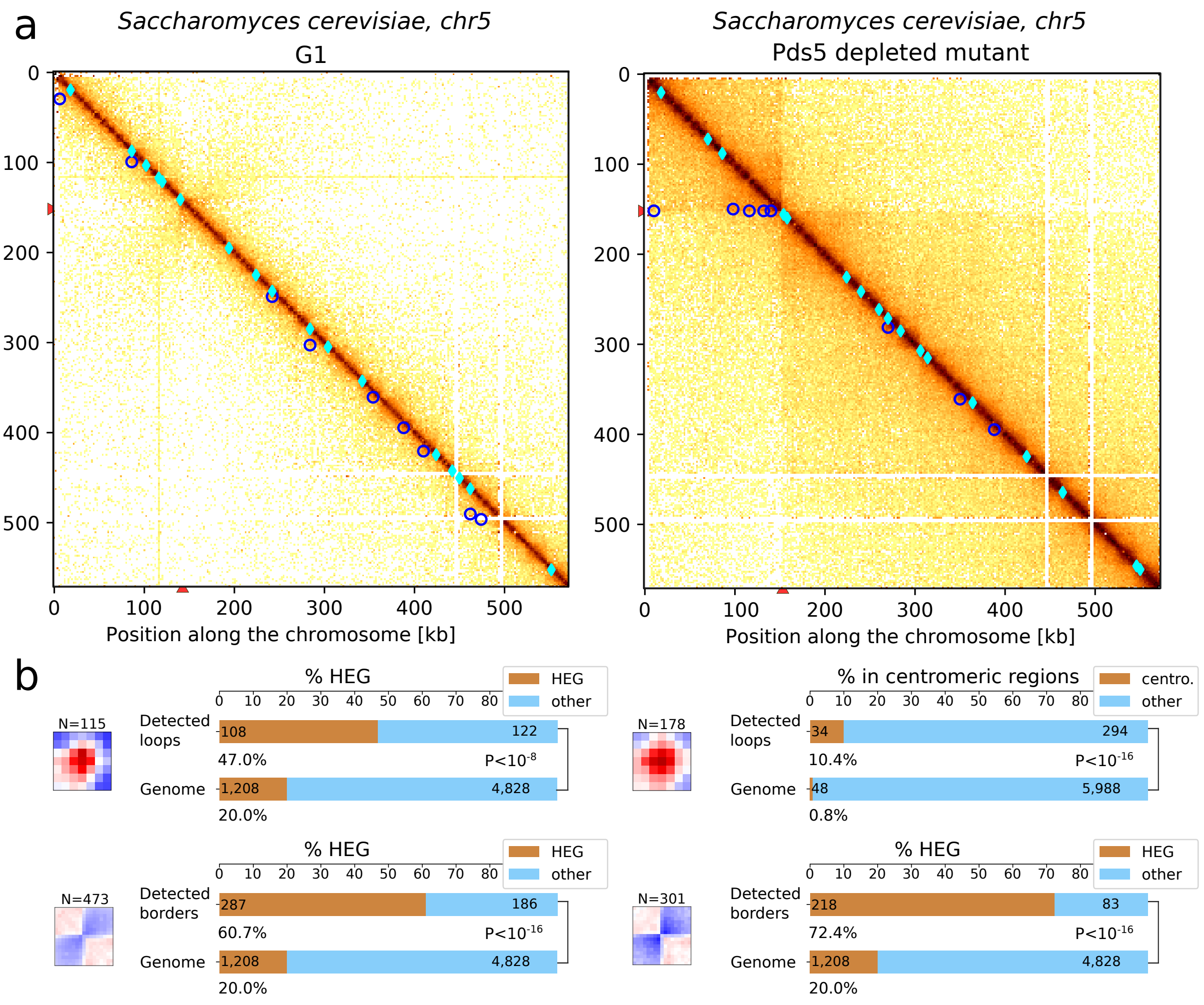


**Fig. 4: Detection of loop and border patterns in yeast contact data a,** Detection of loops and borders in Hi-C data of *S.Cerevisiae* synchronised in G1 and for a mutant depleted in the protein Pds5. **b,** Bar plots showing enrichment in highly expressed genes (HEG) for detected loops in G1 and an enrichment in centromeric regions for the Pds5 mutant. Bar plots showing enrichment in highly expressed genes for detected borders in G1 and Pds5 mutant.

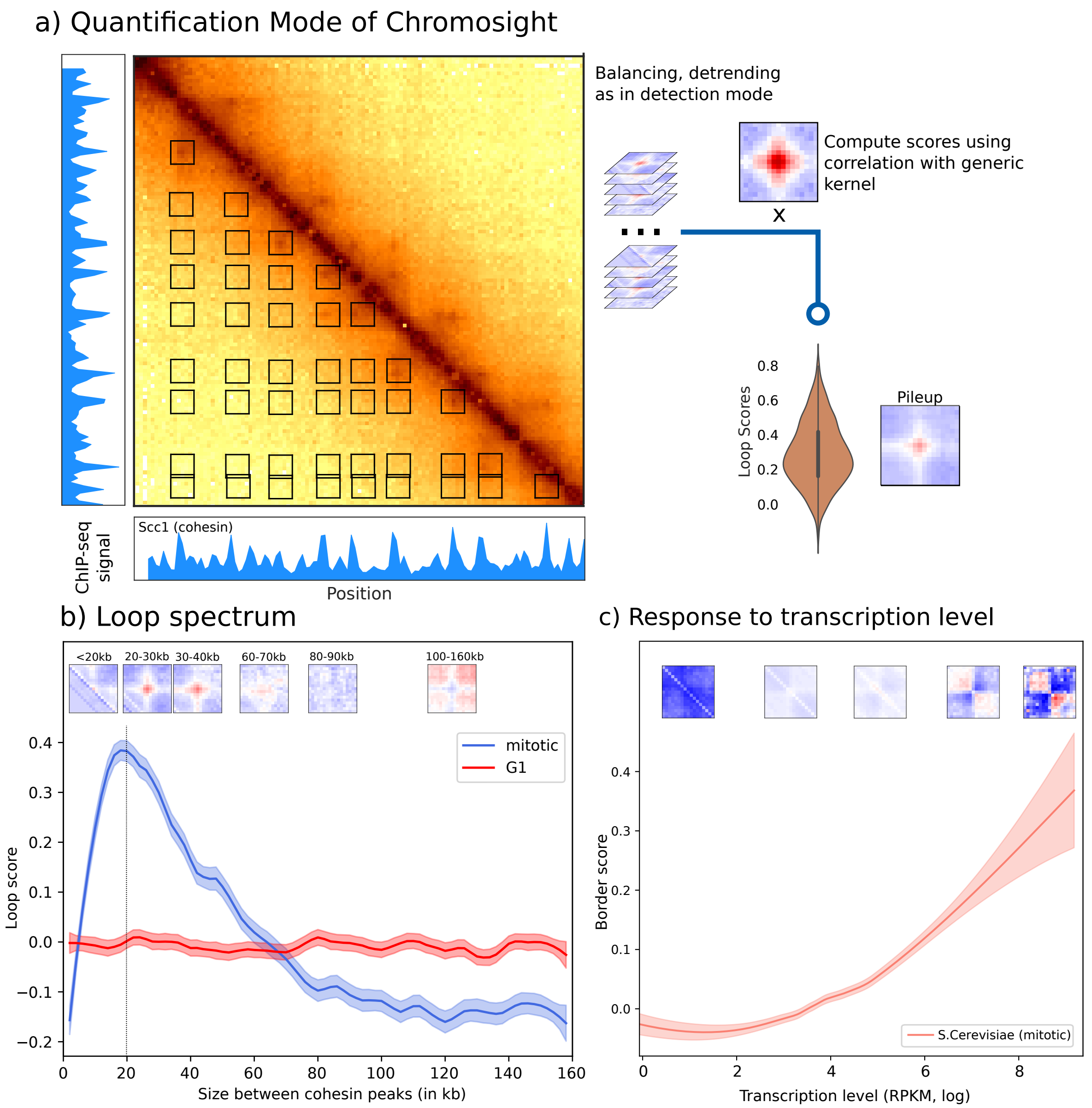


**Fig. 5: Applications of quantification mode on yeast contact data a,** Chromosight quantification mode workflow: sub-matrices from certain 2D genomics positions are extracted from balanced and detrended matrices (as in detection mode). Correlation is then computed for each sub-matrice and the mean of all the sub-matrices is giving a pileup visualisation. 2D genomics positions can pairs of peaks of protein of interest from a ChIP-seq experiement. **b,** Loop spectrum computed for the cohesin loop network. The loop score is given as a function of distance between cohesin peaks for cells in mitotic state (data from [5]). **c,** Plot showing the border score as a funtion of transcription levels in *S.Cerevisiae*, (contact data and transcriptome data from [6]).

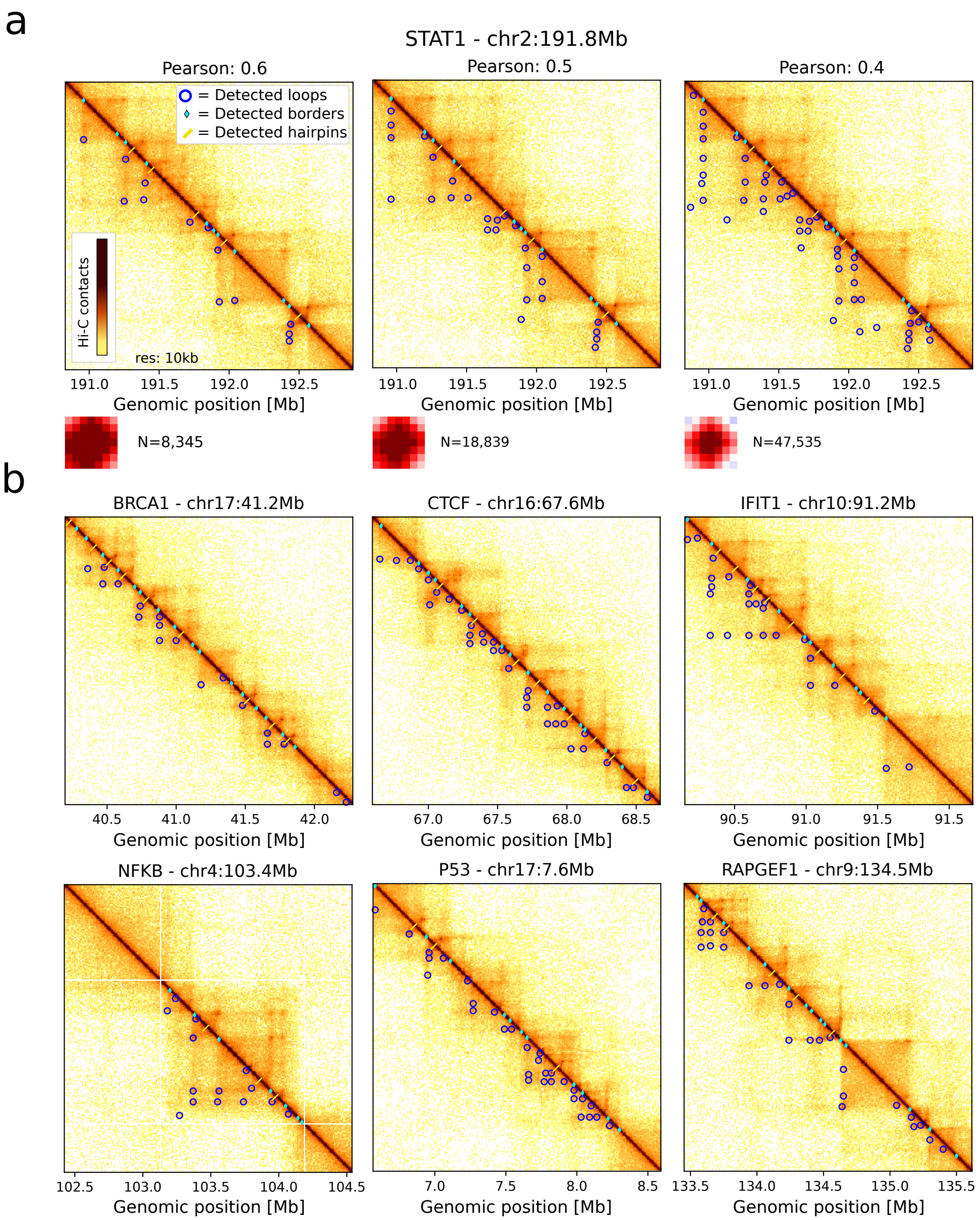


**Fig. 6: Detection of loops in the vicinity of well-studied genes. a,** Effect of decreasing the Pearson coefficient for the loop detection. Contact map in the vicinity of STAT1 gene and total number of detected loops are shown for 3 Pearson coefficients: 0.6, 0.5, 0.4 is shown. Decreasing the Pearson coefficient allows the detection of weak patterns. **b,** Zoom of contact maps 2 Mb around different genes of interest: BRCA1, CTCF, IFIT1, NF$\kappa$B, P53, RAPGEF1 in Hi-C contact data of lymphoblastoids [7]. Dectection done with Pearson coefficient parameter set to 0.5.

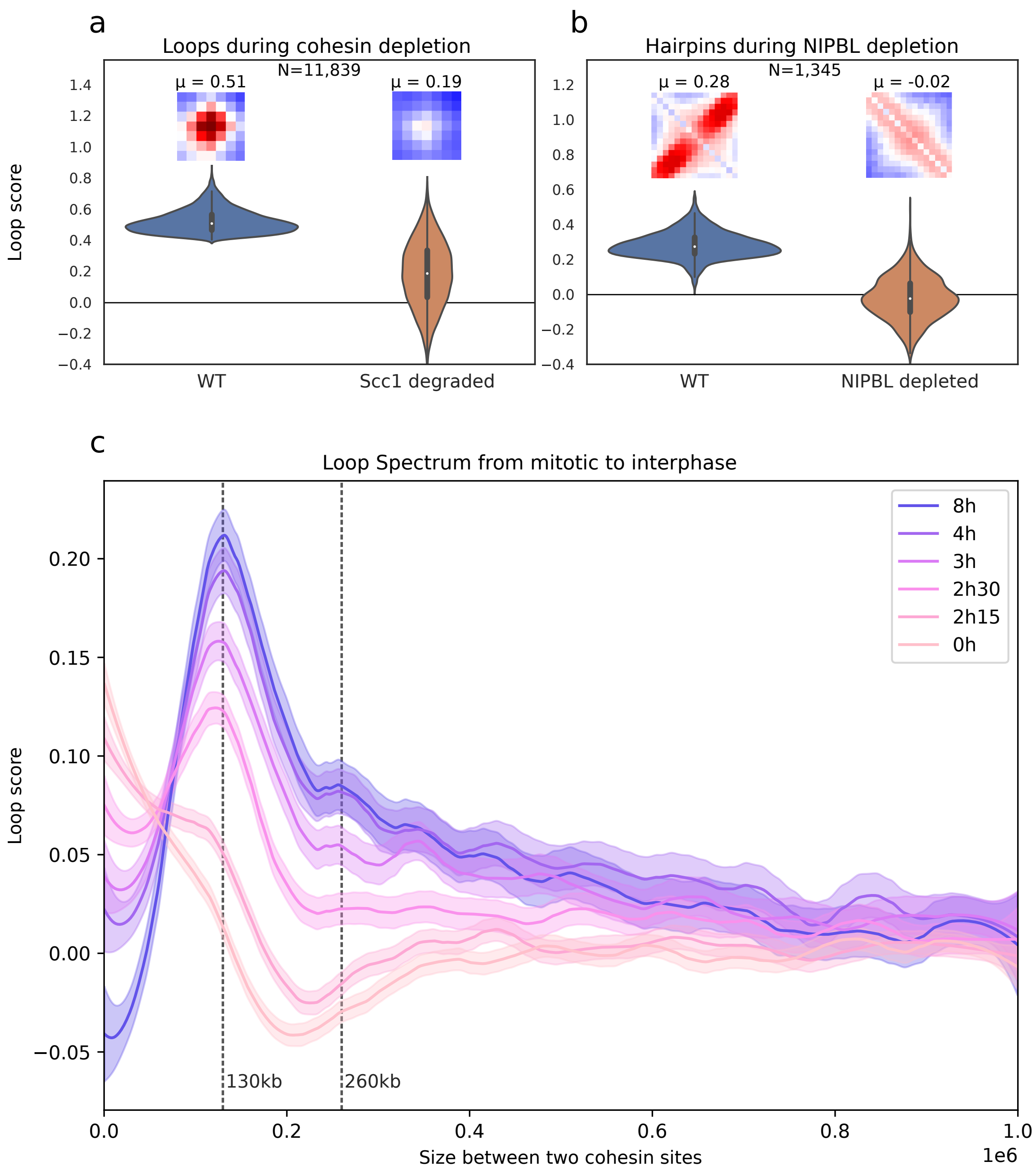


**Fig. 7: Applications of quantification of loops and hairpins on human contact data**. **a,** Comparison of loop score distributions in WT (*Homo sapiens*) and in mutant cells depleted in Scc1 [8] for loops detected in WT condition. Associated pileup plots of windows centered on detected loops in WT condition. $\mu$: median of loop scores. **b,** Comparison of hairpin score distributions in WT (*Mus musculus*, liver cells) and in mutant cells depleted in NIPBL [9] for hairpins detected in the WT condition. Associated pileup plots of windows centered on detected hairpins in WT condition. **c,** Loop spectrum of the loop scores for pairs of Rad21 ChIP-seq peaks separated by increasing distances, at different time points during release from mitosis into G1 [10].

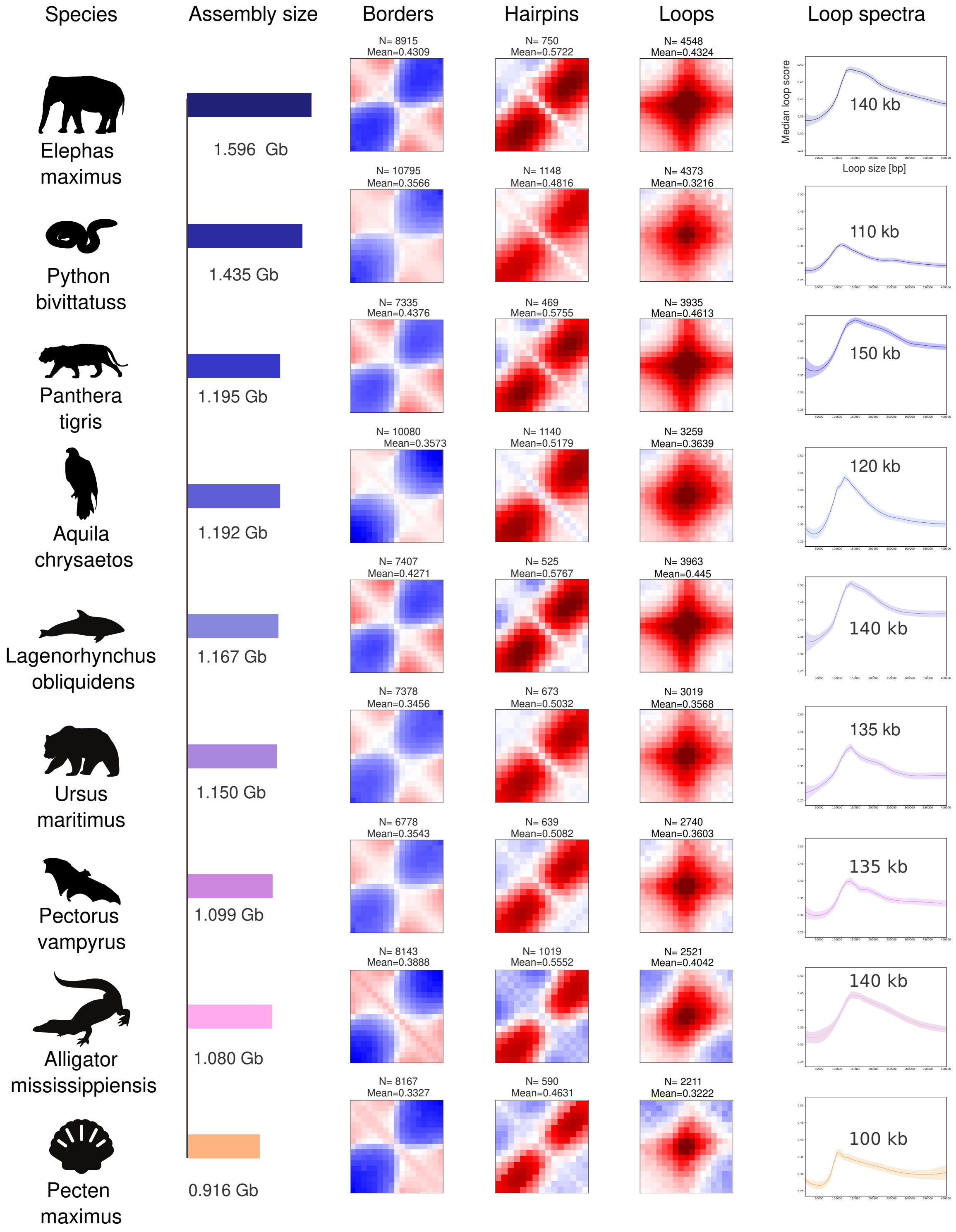


**Fig. 8: Detection of loops, borders, hairpins in various animals from the DNA Zoo project [11].** From left to right, name of the species, barplot giving the genome assembly size, associated pileup plots for detected borders, hairpins and loops patterns and loop spectra computed on the positions of detected loops. The loop spectrum gives the size at which the detected loops have the highest scores. Detection has been performed on a standard laptop with a calculation time of less than 5 min for each pattern per organism.

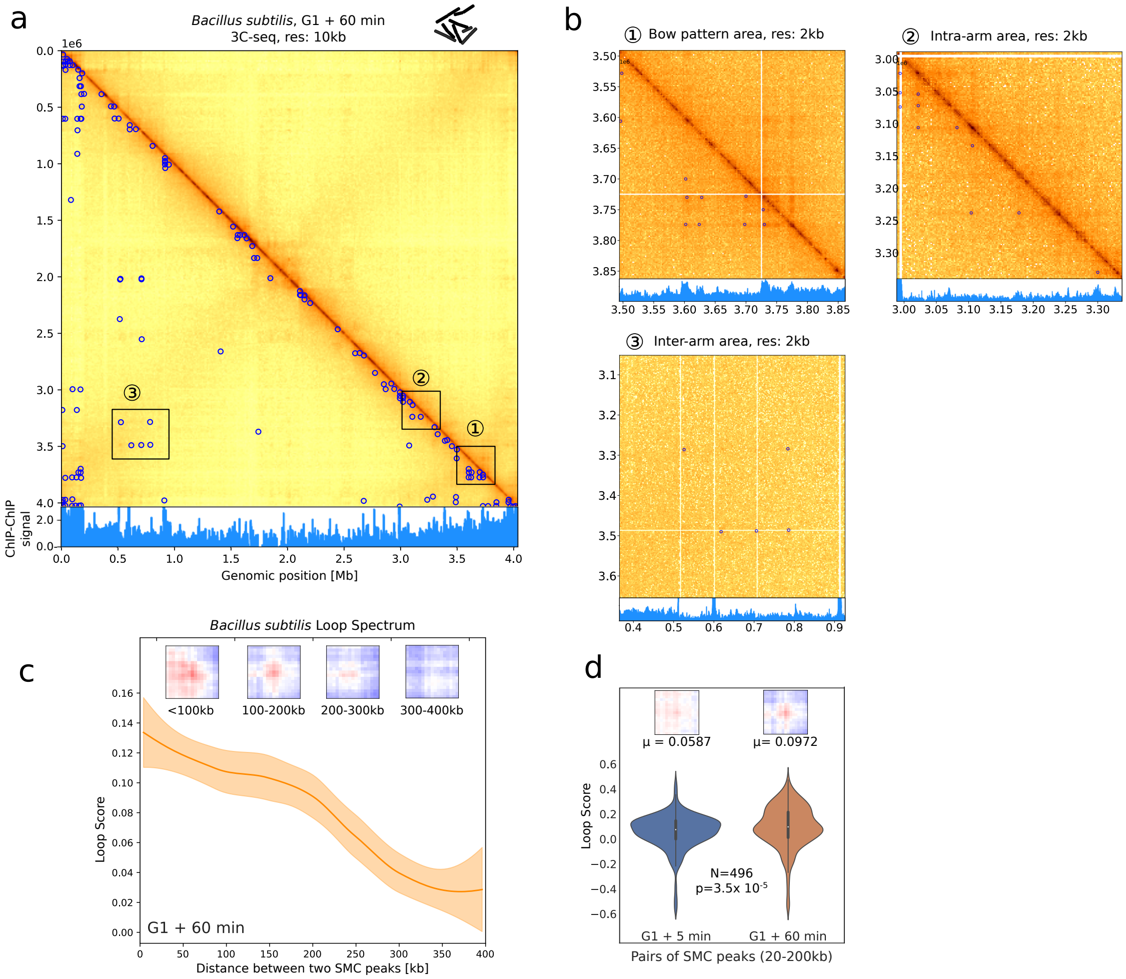


**Fig. 9: Detection and quantification of loops in 3Cseq data of Bacillus subtilis.**
**a,** Detection of loops in 3Cseq data of *Bacillus subtilis* [12]. Genome contact map is shown at 10 kb resolution annotated with detected loops (carried on 2 kb data, 17x17 loop kernel). ChIP-chip signal of Structural Maintenance of Chromosomes proteins (SMC) is plotted under the map. **b,** Zooms of 3 genomic regions highlighted in a): in the bow pattern, in intra-arm region or in inter-arms area.
**c,** Quantification of loop signal for pairs of SMC peaks for different sizes. Associated pileups of patterns for 4 size ranges are shown above. **d,** Quantification of loop signals for pairs of SMC peaks between 20 and 200 kb in 2 conditions: G1 + 5 min and G1 + 60 min. Mean of loop scores and associated p-value (Paired Mann Withney U test).

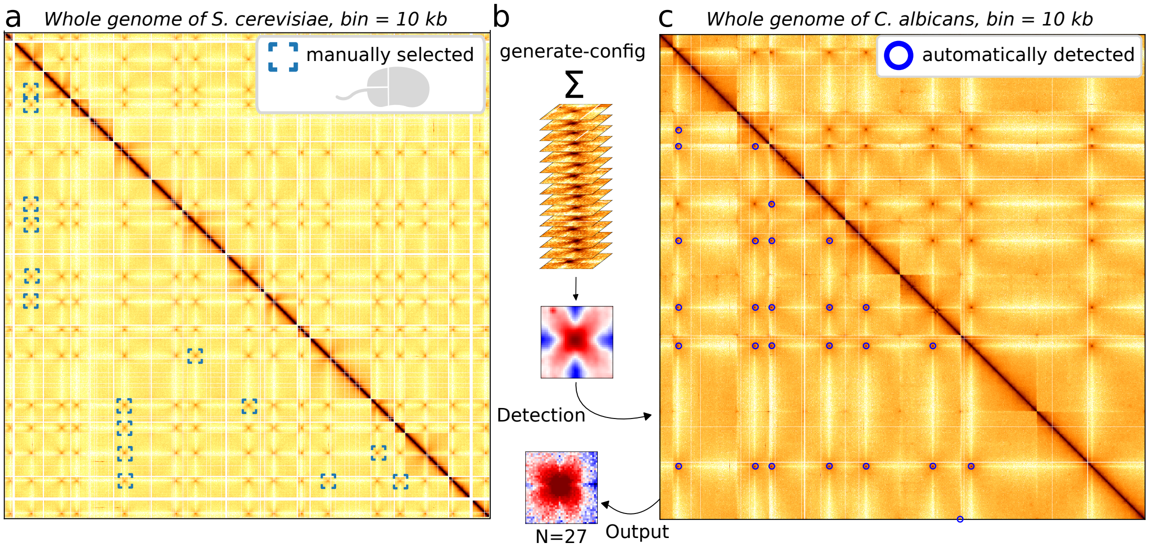


**Fig. 10: Click and Find mode. a,** Whole genome contact map of *S. cerevisiae* [6] with 15 inter-centromere patterns that were selected by hand. **b,** Chromosight generates a new kernel by summing all the selected patterns and applying a Gaussian filter. **c,** Chromosight detection of the inter-centromeres patterns in the whole genome contact map of *C. albicans* [13] with the resulting pileup plot of the 27 detections.

In addition to the kernels presented here (loop, border, hairpin), direct visualisation of the contact maps may lead to the identification of new patterns of interest. We anticipate that many more patterns will be added to the catalogue of visual patterns linked to different molecular mechanisms of chromosome architecture. We have therefore integrated into Chromosight a "Click and Find" mode that allows the user to select patterns that he identifies directly on the contact maps. A new kernel is generated by summing all the selected windows and applying a Gaussian filter to attenuate fluctuations in the case of a small number of selected windows. This kernel can be used in the other modes of Chromosight (detection, quantification) for further analyses. We illustrate this functionality on the centromere-centromere interaction pattern. This cross-shaped pattern is characteristic of the so-called Rabl configuration of genome where the chromosomes are attached to their centromeres [14]. We selected (by double-clicking) 15 patterns of centromere-centromere interactions in the yeast *S.Cerevisiae* and then performed the detection in another species of yeast *Candida Albicans* (at resolution of 5 kb, [13]). Chromosight detected 26 out of the 28 inter-centromeric patterns with one false positive (likely genome misassemblie, located at the edge of the map), which is sufficient to recover the genomic coordinates of all 8 centromeres (**Supplementary Fig 10**). It is also important to note that Chromosight has been able to distinguish patterns of interactions between centrometers from patterns of interactions between telomeres that are also present in the data and that share several common geometrical characteristics. This analysis shows the ability of Chromosight to quickly detect any type of user-defined pattern.

| software | parameter | values |
|---|---|---|
| chromosight | –window-size | 10,15,20 |
| chromosight | –min-dist | 0,40000 |
| chromosight | –pearson | 0.30,0.35,0.40,0.45,0.50 |
| chromosight | –min-separation | 0,50000 |
| chromosight | –full | On,off |
| hicexplorer | –windowSize | 10,15,20 |
| hicexplorer | –peakWidth | 4,5,6,7,8 |
| hicexplorer | –peakInteractionsThreshold | 10,20,30 |
| hicexplorer | –pValuePreselection | 0.01,0.02,0.05,0.1 |
| cooltools | –max-loci-separation | 100000,200000,1000000,2000000 |
| cooltools | –max-nans-tolerated | 5,10,15,20 |
| cooltools | –dots-clustering-radius | 14000,19000,34000,39000 |
| hiccups | -p | 1,2,4,6 |
| hiccups | -i | 6,10,14 |
| hiccups | -f | 0.05,0.1,0.2 |
| homer | -poissonLoopGlobalBg | 0.0001,0.001 |
| homer | -poissonLoopLocalBg | 0.01,0.05,0.1 |
| homer | -window | 2000,5000,10000 |

**Table 1: Parameters used in the benchmark.** Name and value of all parameters used in the benchmark for each software.

| Organism | Experiment type | Figure | Ref | Identifier |
|---|---|---|---|---|
| *S.cerevisiae* | Hi-C, mitotic (nocodazole synchr.) | **Fig** 2 | [6] | SRR7706226, SRR7706227 |
| *S.cerevisiae* | Hi-C, G1 (alpha factor synchr.) | **Fig** 2 | [5] | SRR8769554 |
| s *S.cerevisiae* | Hi-C, Scc1 degron, mitotic (cdc20 synchr.) | **Fig** 2 | [5] | SRR8769548 |
| *S.cerevisiae* | Hi-C, meiosis (t=0h, 3h, 4h 6h) | **Sup Fig** 4 | [15] | SRR7126297 SRR7126293 SRR7126301 SRR7340033 |
| *S.cerevisiae* | Hi-C, Pds5 depleted, mitotic (cdc20 synchr.) | **Sup Fig** 4 | [5] | SRR8769553 |
| *S.pombe* | Hi-C, Mitotic phase, 40 min | **Fig** 2 | [16] | SRR5149256 |
| *H.sapiens* | Hi-C, GM12878, asynchronous | **Fig** 3 | [7] | SRR6675327 |
| *H.sapiens* | Hi-C, Hela cells, WT | **Fig** 3 | [8] | GSM2747745 |
| *H.sapiens* | Hi-C, Hela cells, depleted in Scc1 | **Fig** 3 | [8] | GSM2747747 |
| *H.sapiens* | Hi-C, Hela cells during cell cycle (R2, T0, T2h15, T2h30, T3h, T4h, T8h) | **Fig** 3 | [10] | GSM3909703 GSM3909697 GSM3909696 GSM3909694 GSM3909691 GSM3909686 |
| *M.musculus* | Hi-C, liver cells | **Fig** 3 | [9] | GSE93431 |
| *M.musculus* | Hi-C, liver cells, depleted in NIPBL | **Fig** 3 | [9] | GSE93431 |
| *B.subtilis* | 3Cseq in G1 + 5 min | **Fig** 3 | [2] | SRR2214069 |
| *B.subtilis* | 3Cseq in G1 + 60 min | **Fig** 3 | [2] | SRR2214080 |
| *Epstein Barr Virus* | ChIA-PET of CTCF in GM12878 cells | **Fig** 3 | [17] | SRR2312566 |
| *C. Albicans* | Hi-C, asynchronous | **Fig** 10 | [13] | SRR3381672 |
| *E. maximus* | Hi-C, asynchronous | **Sup Fig** 8 | [11] | Elephas_maximus rawchrom.hic |
| *P. bivittatuss* | Hi-C, asynchronous | **Sup Fig** 8 | [18] | Python_bivittatuss rawchrom.hic |
| *P. tigris* | Hi-C, asynchronous | **Sup Fig** 8 | [19] | Panthera_tigris rawchrom.hic |
| *A. chrysaetos* | Hi-C, asynchronous | **Sup Fig** 8 | [20] | Aquila_chrysaetos rawchrom.hic |
| *L. obliquidens* | Hi-C, asynchronous | **Sup Fig** 8 | [11] | Lagenorhynchus _obliquidens rawchrom.hic |
| *U. maritimus* | Hi-C, asynchronous | **Sup Fig** 8 | [21] | Ursus_maritimus rawchrom.hic |
| *P. vampyrus* | Hi-C, asynchronous | **Sup Fig** 8 | [22] | Pectorus_vampyrus rawchrom.hic |
| *A. mississippiensis* | Hi-C, asynchronous | **Sup Fig** 8 | [23][24] | Alligator_mississi -ppiensis rawchrom.hic |
| *P. maximus* | Hi-C, asynchronous | **Sup Fig** 8 | [25] | Pecten_maximus rawchrom.hic |

**Table 2: Different contact datasets analysed in the present study.** The last column indicates either the identifier for the raw reads available on the Short Read Archive server (SRA) (https://www.ncbi.nlm.nih.gov/sra), the identifier of the .cool files accessible on the Gene Expression Omnibus server (GEO) https://www.ncbi.nlm.nih.gov/geo or the name of hic files from DNA zoo project available on https://www.dnazoo.org/assemblies [11] from which the analysis were made.

| Organism | Experiment type | Figure | Ref | Identifier |
|---|---|---|---|---|
| *S.cerevisiae* | RNA-seq, mitotic (nocodazole synchr.) | **Fig** 2 | [6] | SRR7692240 |
| *S.cerevisiae* | ChIP-seq, Scc1PK9 IP G1 releasing 60min | **Fig** 2 | [26] | SRR2065097, SRR2065092 |
| *H.sapiens* | ChIP-seq CTCF | **Fig** 3 | [27] | wgEncodeAwgTfbsBroad Gm12878CtcfUniPk.narrowPeak |
| *H.sapiens* | ChIP-seq RAD21 | **Fig** 3 | [27] | wgEncodeAwgTfbsHaib Gm12878Rad21V0416101UniPk |
| *H.sapiens* | ChIP-seq NIPBL | **Fig** 3 | [27] | GSM2443453_GM12878_NIPBL Rep1_ 2WCE_Narrow_Peaks_peaks.narrowPeak |
| *B.subtilis* | ChIP-chip of SMC | **Fig** 3 | [28] | GSE14693 |
| *Epstein Barr Virus* | ChIP-seq CTCF | **Fig** 3 | [29] | SRR036682 |
| *Epstein Barr Virus* | ChIP-seq RAD21 | **Fig** 3 | [17] | SRR2312570 |

**Table 3: Other genomic datasets used in the present study.** The last column indicates either the identifier for the raw reads available on the Short Read Archive server (SRA) (https://www.ncbi.nlm.nih.gov/sra), the identifier of the ChIP-chip files accessible on the Gene Expression Omnibus server (GEO) https://www.ncbi.nlm.nih.gov/geo or the identifier of ChIP-seq peak files available on http://genome.ucsc.edu.
